# A High-Content Pseudotime Framework for Morphological Profiling of Dengue Virus Inhibitors from a Drug-Repurposing Library

**DOI:** 10.64898/2026.10.06.757061

**Authors:** Jillian G. Hoffstadt, Samantha F. Boisvert, Jesse W. Wotring, Samuel Porter, Benjamin Halligan, Daniel Macedo de Melo Jorge, Andrew W. Tai, Jonathan Z. Sexton, Matthew J. O’Meara

## Abstract

Dengue virus (DENV) causes dengue fever, a globally prevalent, epidemic-prone disease with no FDA-approved antivirals. Additionally, the available vaccine for DENV can increase the risk of severe dengue fever for those who have never had a DENV infection due to antibody-dependent enhancement. Thus, there is an urgent need to identify DENV antivirals. Drug repurposing, high-content screening, and machine learning were leveraged to efficiently identify antivirals likely to inhibit DENV replication. Using high-content screening in a DENV infection model, we characterized the morphological patterns of NS4B and envelope (E) protein expression over time and developed a viral pseudotime model that predicts infection progression to aid in phenotypic interpretation of drug effects. We then developed a single cell infection classifier for antiviral efficacy and performed high-throughput drug screening of 960 compounds. We identified four concentration-dependent inhibitors of DENV with nanomolar-to-submicromolar potencies: esomeprazole, GW4064, pralatrexate, and LY411575. With our pseudotime machine learning model trained on data from the time-series assay, we scored the concentration-dependent inhibitors to map their mode of action onto time points in the viral life cycle. Our most notable results show that LY411575, a gamma secretase inhibitor, exhibited an IC_50_ of 72 nM and reduced percent infection to near the level of the mock infection control. Pseudotime inferences generated mode of action predictions that were supported by time-of-addition experiments, indicating that LY411575 arrests DENV infection before the development of late-stage infection associated morphologies, while retaining activity when added after inoculation.

**Importance:** Our approach reported herein couples high-content screening and a viral pseudotime model as a robust platform for the identification of viral inhibitors with imputation of morphological profiles for large-scale drug discovery efforts. Viral pseudotime models can, in principle, be developed for any viruses or biological processes through a defined life cycle, suggesting that phenotypic time points, or phases, can be mapped across data in a broad range of applications. We report a high-content pseudotime framework for mapping temporal phenotypes and apply it to drug repurposing for DENV antivirals, gaining insight into how the mode of action of the screened antivirals correlates to phases of DENV replication. This study demonstrates how image-derived phenotypes can be leveraged to generate mode of action hypotheses and prioritize antiviral candidates for follow-up studies.

## Introduction

The global risk of dengue fever has grown dramatically, with an estimated 100-400 million annual infections and 96 million symptomatic cases,^1^ ranging from mild flu-like cases to dengue shock syndrome (DSS) or dengue hemorrhagic fever (DHF) characterized by severe bleeding, organ failure, and plasma leakage that can be fatal.^2^ Rising temperatures are expanding the mosquito’s vector range to new regions, including Europe and the Eastern Mediterranean.^1^

Dengue fever is caused by dengue virus (DENV), a positive-sense single-stranded RNA virus in the Flaviviridae family with four serotypes. Recovery from one serotype confers lifelong immunity to it, but subsequent infection with a different serotype increases risk of severe dengue via antibody-dependent enhancement.^3^ The live-attenuated vaccine CYD-TDV carries this same risk in seronegative recipients, so its FDA indication (Dengvaxia) was restricted to ages 6 to 16 with laboratory-confirmed prior infection in endemic areas, but has since been discontinued.^4^ A second vaccine, TAK-003 (Qdenga), is now WHO-recommended for children aged 6 to 16 in high-transmission settings, but restricted indications, serostatus requirements, and uneven availability leave a substantial unmet need for antivirals.

Currently, no FDA-approved small molecule antivirals exist for any DENV serotype. **Table 1** summarizes clinical candidates that have reached trials for DENV, active and discontinued. Small molecules evaluated in these trials have largely proven to be ineffective or were discontinued, the most promising being mosnodenvir (JNJ-1802 or JNJ-64281802), a pan-serotype NS4B inhibitor.^5–11^ Despite encouraging clinical trial results, development of mosnodenvir was discontinued in 2024 following changes in portfolio priorities, with no reported safety or efficacy concerns.^11,12^

**Table 1.** Clinically evaluated compounds for DENV. This table lists small molecules with completed clinical trial data for DENV (current as of August 2026). Ongoing trials without reported results (e.g., dexamethasone, anakinra, zanamivir) and monoclonal antibody candidates are not included.

| Drug Name | Screening Strategy | Activity/Target | Clinical Trial Phase | Status |
| --- | --- | --- | --- | --- |
| **Balapiravir**<sup>5</sup> | *In vitro* | Polymerase inhibitor | Phase 1<br>(NCT01096576) | Ineffective |
| **AT-752**<sup>6</sup> | *In vitro* and *in vivo* | NS5 polymerase;<br>nucleotide prodrug | Phase 2<br>(NCT05466240) | Terminated |
| **NITD-688**<br>**(EYU688)**<sup>7</sup> | *In vitro*<br>Phenotypic | NS4B inhibitor | Phase 2 | Ongoing |
| **Celgosivir**<sup>8</sup> | *In vitro*/<br>immunodetection | Alpha-glucosidase I<br>inhibitor | Phase 1b<br>(NCT01619969) | No<br>significant<br>reduction in<br>viral load or<br>fever |
| **Celgosivir/**<br>**Modipafant** | *In vitro* | Celgosivir: alpha-glucosidase I<br>inhibitor;<br>Modipafant: PAF<br>receptor antagonist | Phase 1/2<br>(NCT02569827) | Withdrawn<br>(0 enrolled) |
| **Montelukast**<sup>9</sup> | FDA approved<br>(1998); *in vivo*<br>Randomized<br>Controlled Trial | Leukotriene<br>receptor antagonist | Phase 2/3<br>(NCT04673422) | Ineffective |
| **Molnupiravir<sup>10</sup>** | *NA* | Induce mutational load | Viral clearance study (NCT06551844) | Ongoing |
| **Ivermectin<sup>10</sup>** | Repurposed antiparasitic | Activates glutamate-gated chloride channels | Phase 2/3 (NCT02045069) | Ineffective |
| **Chloroquine<sup>10</sup>** | Repurposed antiparasitic | Proposed antiviral/anti-inflammatory activity | Phase 1/2 (NCT00849602) | Ineffective |
| **Mosnodenvir (JNJ-1802)<sup>11</sup>** | *In vitro* | Inhibits NS3-NS4B complex | Phase 2 (NCT04906980) | Terminated |

A promising strategy for antiviral discovery with potential for rapid deployment is drug repurposing, using compounds with established safety profiles for new indications. Repurposing libraries are annotated with targets and mechanisms, so pairing this information with morphological phenotype allows informed hypotheses about a hit compound’s mechanism of action, but this requires a screening readout that captures not just whether a compound blocks infection, but how it perturbs the cell.

Phenotypic screening is a powerful approach for identifying small molecule inhibitors across disease areas. High-content imaging, a form of phenotypic screening, provides high-dimensional data via automated multi-fluorescent microscopy and high-throughput computational analysis. Unlike conventional screens limited to a few hand-picked endpoints, high-content imaging draws on hundreds to thousands of cellular features,^13^ well suited to characterizing DENV infection dynamics and antiviral candidates.^14^ Because these features vary continuously as infection progresses, morphology can map cells onto their position in the infection cycle.

Phenotypic screens have delivered promising leads for DENV and other related viruses.^15–18^ For example, a phenotypic anti-dengue screen with an eGFP readout (KU Leuven in partnership with Janssen Pharmaceutical) identified JNJ-A07 as a potent inhibitor of NS3-NS4B protein interaction in VeroE6 cells,^15^ illustrating how target-agnostic screens can reveal therapeutic targets, like NS4B, without prior bias. Viral proteins lacking host homologs, like NS4B, are attractive targets for potentially greater selectivity and reduced off-target risk. For this study, NS4B and E protein where stained for in Huh7.5.1 cells, aiming to identify morphological endpoints of NS4B antagonists or other DENV inhibition mechanisms.

In this study, we hypothesize that combining high-content imaging and drug repurposing will identify DENV inhibitors and provide valuable insight into their modes of action. To more precisely screen for DENV antivirals, we first aimed to understand the morphology of the DENV infection trajectory in Huh7.5.1 cells by characterizing the detection and spatial localization of NS4B and the envelope (E) protein in a pseudotime assay over 48 hours in the absence of drug treatment. Then, through high-content phenotypic screening, we screened a repurposing library and used machine learning to generate hypotheses for hit compounds’ mechanism of action.

Using the morphology information captured in the pseudotime assay, we can infer a compound’s mode of action by imputing its pseudotime of arrest, the point along the infection cycle at which it arrests the cell. Herein, we describe a DENV discovery platform that delivers a novel technique for morphological profiling and, to our knowledge, for the first time, the mapping of DENV inhibitor mechanisms directly onto stages of the viral infection cycle.

## Methods

### Compounds

The University of Michigan Drug Repurposing Library (DRL) was assembled in the Sexton laboratory from Selleck Chemicals and Cayman Chemical and prepared as 2 mM stock solutions in dimethylsulfoxide (DMSO) in Echo PP Plates. 960 compounds from the DRL were dispensed using a Beckman Coulter Echo 650 acoustic liquid handler via a 100 nL transfer to cell plates, giving a nominal final screening concentration of 4 μM in 0.2% DMSO after addition of the cell suspension; the same dispensing procedure was used for concentration-response testing. For time-of-addition experiments, four concentration-responsive screening hits (esomeprazole, pralatrexate, LY411575, and GW4064) were repurchased.

### Cells and Virus

The human hepatoma-derived Huh7.5.1 cells were maintained in Dulbecco’s Modified Eagle Medium (DMEM) supplemented with 10% fetal bovine serum (FBS) and 1X penicillin-streptomycin solution (15140122, Gibco) and grown at 37 °C with 5% CO2 following standard cell culture procedures. To generate wild-type DENV-2 virus, the pD2/IC-30P-NBX plasmid was linearized with XbaI restriction enzyme digestion, in vitro transcribed, and capped with the m7G(5′)ppp(5′)A cap analog using T7 Megascript; this RNA was transfected into Vero E6 cells using TransIT mRNA, and virus stock titers were determined by TCID50 using the Reed and Muench method. All work with live DENV virus was performed at biosafety level 2 (BSL2) with the approval of the University of Michigan’s Department of Environment and Health and Safety and the Institutional Biosafety Committee.

### Anti-DENV High-Content Bioassays

For drug screening, compounds were dispensed as described above. Huh7.5.1 cells were seeded on top of the compounds at 3,000 cells per well in 50 μL of media. After 24 hours of cell attachment and compound pre-incubation at 37 °C and 5% CO_2_, cells were inoculated with DENV at a multiplicity of infection (MOI) of 2. Cells were incubated with virus and compounds for an additional 48 h and then fixed with 100% methanol for 10 mins at −20 °C. Cells were then permeabilized with 0.3% Triton-X100 for 20 minutes. The plates were then incubated with an anti-NS4B primary antibody (GTX103349, GeneTex, Irvine, California, USA) at a dilution of 1:2000 overnight at 4 °C. The staining buffer was PBS with 1.5% w/v BSA flakes, 1% v/v goat serum, and 1X TBS Tween 20. Following primary antibody staining, cells were then stained with 1:1000 secondary antibody Alexa Fluor 647 for NS4B labeling (A-31573, donkey anti-rabbit, Thermo Fisher, Waltham, MA, USA), 1:4000 HCS CellMask Orange (H32713, Thermo Fisher, Waltham, MA, USA) for cytoplasmic labeling, 1:25 Concanavalin A (C11252, Thermo Fisher, Waltham, MA, USA) for ER labeling, and 1:1000 Hoechst 33342 (H1399, Thermo Fisher, Waltham, MA, USA) for nuclear labeling. All dyes were incubated for a total of 45 mins at room temperature. Cells were washed and stored in PBS before imaging.

For the pseudotime series, Huh7.5.1 cells were seeded as described above. After 24 hours of cell attachment at 37 °C and 5% CO_2_, cells were inoculated with DENV (MOI=2) and subsequent inoculations were performed after a 12 h interval and then at 8 h intervals, giving 48, 36, 28, 20, 12, and 0 hours post infection (hpi) conditions, with each time point occupying a unique four-column block of 64 wells within the same plate. The 0 hpi condition was performed as a mock infection where no virus was added as a control. All cells were fixed at the same time and stained following the same procedure as the drug screen but using anti-NS4B protein primary antibody at a dilution of 1:2000 and a 4G2 anti-E protein primary antibody produced in-house^18^ at a dilution of 1:10 overnight at 4 °C. Following primary antibody staining, cells were stained with 1:1000 secondary antibody Alexa Fluor 647, 1:4000 HCS CellMask Orange, 1:1000 secondary antibody Alexa Fluor 488 for E protein labeling (A-21121, goat anti-mouse, Thermo Fisher, Waltham, MA, USA), and 1:1000 Hoechst 33342.

For the follow-up concentration-response assay, plates were treated as described for the drug screen, except compounds were dispensed in a 10-point, 3-fold serial dilution starting from 8 μM. Concentration-response confirmation was planned for all eight primary screening hits, but one hit, buflomedil HCl, could not be tested due to a compound-dispensing failure. Two additional compounds, remdesivir and Z-FA-FMK, were included in the concentration-response experiment based on the laboratory’s concurrent interest in another antiviral study. Concentration-response data are therefore reported for seven of eight primary hits plus two investigational compounds (n=9 total).

### High-Content Imaging

Stained assay plates were imaged using a Yokogawa Cell Voyager 8000 (CV8000) microscope with 20X/1.0NA water immersion lens. Maximum intensity projection images were collected over a 10 μm Z-height with 2 μm spacing. A total of 9 fields per well were imaged for all assay plates, accounting for roughly 80% of the total well area. Laser power and camera exposure times were adjusted to yield optimal signal to noise.

### Image Processing

Nuclear and cell segmentation was performed using the CellPose 2.0 segmentation software^19^ (diameter=40, no edge, model=cyto 2) and CellProfiler 4.0.^20^ CellProfiler was used for background subtraction and feature extraction to measure per-cell viral protein intensity and texture features (Alexa Fluor 647 & 488) as well as general cell morphologic measurements. 342 features (Texture, Intensities, Cell Shape, and Area) were collected for each cell. Further details on the image-processing pipeline are provided in the Data Availability section. A random forest^21^ cell infection classifier was trained on approximately 100 infected and 100 uninfected cells by manual selection using the CellProfiler Analyst 3.0.4,^22^ achieving 99.5% accuracy under 5-fold cross-validation.

### Time-Series Analysis

Cell-level features of 0 and 48 hpi for the NS4B and E protein (“NS4B & E pro” model) channels were used to train XGBoost^23^ XGBRegressor estimators with the default squared-error regression objective to score infection under 5-fold cross-validation performed using Scikit-learn.^24^ Only fields with cell counts above 200 were included in the training set. To model infection dynamics, separate models for only NS4B (“NS4B only”) and only E protein (“Epro only”) were trained independently. All cells from the pseudotime series assay were then scored for infection by each of the three models and averaged to the image (field) level and the well level. Scores of infection were compared for statistical significance in a paired t-test.^25^

Cellular features for the NS4B and E protein image channels were also used in a dimensionality reduction analysis using Uniform Manifold Approximation and Projection (UMAP)^26^ for X and Y coordinates. Due to the large number of cell observations, a 20% random subset (650,000 cells) was analyzed.

### Drug Treatments vs. Infection Time Points Analysis

Cell-level features from the pseudotime series assay for 0, 12, 20, 28, 36, and 48 hpi were added to the training data for additional XGBoost regression models as described in the prior section. Each time point was replicated in 64 wells, but after filtering for cell count, 44 wells for hour 0 were included, 53 wells for hour 12, 24 wells for hour 20, 22 wells for hour 28, 18 wells for hour 36, and 44 wells for hour 48. The positive control and negative control wells from the drug treatment experiment were grouped with the 0 and 48 hpi wells, respectively, from the time points analysis training data. Plate-based batch correction was performed using Harmony.^27^ All drug treated wells were subsetted as the scoring data. The XGBoost regression models were trained pairwise for each time point (ex. 12 hpi vs. 36 hpi) with the earlier time point being labeled as 0 and the later time point as 1. Additional XGBoost regression models were trained on one time point versus all others (ex. 12 hpi vs. 0, 20, 28, 36, and 48 hpi) in which the single time point was labeled 0 and all other time points were labeled as 1. The resulting 21 models were all trained under 5-fold cross-validation performed using Scikit-learn. An overview of model training is outlined in **Supplementary Figure S1** and R^2^ values found in **Supplementary Table S1.**

We scored the drug-treated wells of the concentration-response experiment to infer the pseudotime of arrest, reasoning that a confident score under a discriminative model is evidence that the arrested phenotype lies near one of that model’s anchor timepoints. Let *t* ∈ [0, 48] h denote the candidate time of arrest and let *f*(*t*; *μ*) denote a Student’s *t* density with location *μ*, scale *σ* = 6 h, and *ν* = 2 degrees of freedom; the heavy tails at *ν* = 2 make the inference robust to individual mis-scoring models. For a model trained to discriminate time points *t_i_* < *t_j_*, with the earlier time point labeled 0 and later time point labeled 1, we took the observed score *s_ij_* ∈ [0,1] as a mixture weight and defined the per-model likelihood as a two-component mixture, *L_ij_*(*t*|*s_ij_*) = (1 − *s_ij_*) ⋅ *f*(*t*; *t_i_*) + *s_ij_* ⋅ *f*(*t*; *t_j_*). For a model trained to discriminate a single time point *t_j_* (labeled 1) against all other time points (labeled 0), we defined a mixture of the anchored density and a uniform component representing the absence of information when the phenotype did not resemble *t_j_*, *L_j_*(*t*|*s_j_*) = *s_j_* ⋅ *f*(*t*; *t_j_*)+ (1 − *s_j_*) ⋅ 1/48. Both forms are proper densities in *t* for every value of *s*, so models contribute equal total evidence regardless of how confident they are. Because the 21 models are trained on overlapping data and are therefore strongly dependent, we combined them using a logarithmic opinion pool with equal weight *w* = 1/21 rather than by assuming conditional independence: log *L*(*t*|*S*) = *w*[∑*_ij_* log *L_ij_*(*t*|*s_ij_*) + ∑*_j_* log *L_j_*(*t*|*s_j_*) + const. Likelihoods were evaluated on a 0.1 h grid, computed independently for each well, and normalized over [0,48] h to give a proper posterior density under a uniform prior on the time of arrest; densities were then averaged across replicate wells within each compound and concentration. We report a posterior mode and a 95% equal-tailed credible interval. Where the posterior is monotone across the whole window the model falls on a boundary; we report these as unresolved (≤ 0 h or ≥ 48 h) rather than as point estimates. Because XGBoost regression with a squared error objective is unbounded, scores were clipped to [0,1] before use. Sensitivity to *σ* ∈ {3, 6, 9, 12} and *ν* ∈ {1, 2, 4, ∞} is reported in **Supplementary Figure S2.**

### Time-of-Addition Assay

A time-of-addition assay was performed using the same fixation and staining treatments as the concentration-response experiment, omitting the E-pro antibody. The four concentration-responsive compounds (esomeprazole, GW4064, pralatrexate, LY411575) were evaluated at 8 μM (n=14 replicates) at six time points relative to infection: −24 h, −2 h, co-treatment, +2 h, +4 h and +24 h. Huh7.5.1 cells were seeded at 3,000 cells per well in 50 μL of media at 37 °C and 5% CO2, cells were inoculated with DENV at a multiplicity of infection (MOI) of 2. Virus and/ or compound remained on cells continuously from the point of addition until fixation. Cells were fixed and stained at 48hpi. Statistical significance of the time-of-addition assay was assessed within each compound separately by one-way ANOVA followed by Dunnett’s multiple comparisons test, comparing each condition (uninfected control, −24 h, −2 h, co-treatment, +2 h, +4 h, +24 h) to the infected (DMSO) control.

## Results

### Time-Course (Pseudotime) Analysis of DENV Infection

Morphologic profiling of cells infected with DENV at 0, 12, 20, 28, 36, and 48 hpi (**Figure 1A**; Methods) revealed characteristics of the infection cycle. Huh7.5.1 cells were selected for their DENV-permissive cytopathic effect,^28^ and the first time point (12 hpi) approximates the earliest release of infectious progeny from DENV-infected Huh7 cells,^29^ from which Huh7.5.1 cells are derived. Representative images from each stage are shown in **Figure 1B**.

**Figure 1:**
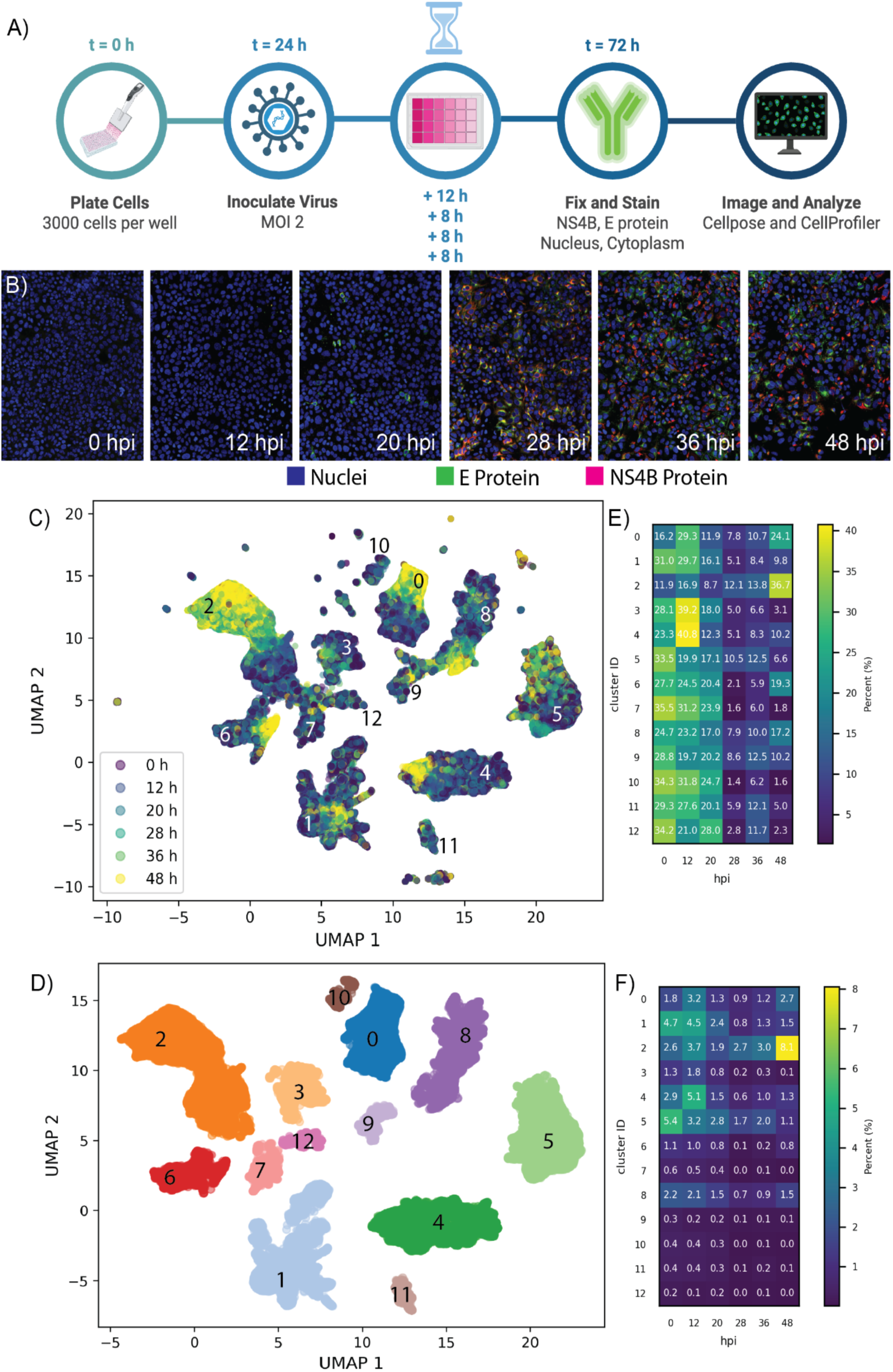
UMAP of cells shows clustering based on time point. A.) Time points infection workflow. B.) Representative images of each hpi in the pseudo-time-series assay. C.) The UMAP embedding of cell-level morphological features shows several distinct clusters of infected cells. The clusters are colored by condition, which is time post infection in hours. D.) A 20 % subset of the data in (C) but colored by cluster ID based on DBSCAN labeling. E.) A heatmap of time post infection versus cluster ID where the percentages are per cluster. Each row (cluster ID) adds up to 100% and shows the distribution of time points per cluster. F.) A heatmap of time post infection versus cluster ID where the percentages are per all cells. The entire heatmap equals 100%.

Intensity, texture, and area/shape measurements were extracted per cell for each time point and analyzed with machine learning. Cell-level measurements from the nuclei, cytoplasm, NS4B, and E protein channels produced a UMAP embedding of a 20% random cell subset (random_state=42; **Figure 1C, D**), showing 13 distinct clusters whose membership was not uniform with respect to time. Coloring by hpi (**Figure 1C**) shows early time points (0–20 hpi) concentrated in the left/lower regions, while 48 hpi cells accumulate mostly in cluster 2 and pockets of clusters 0, 4, 6, and 8. The corresponding DBSCAN^30^ cluster assignments computed with eps = 0.9 and min_samples = 600 are shown in **Figure 1D**.

The heatmap in **Figure 1E** quantifies the UMAP distribution, showing the percentage of each cluster’s cells drawn from each time point. Several clusters are strongly enriched for early infection: clusters 7, 10, and 12 draw most of their cells from 0–20 hpi (90.6%, 90.8%, and 83.2%, respectively), while clusters 3 and 4 peak at 12 hpi (39.2% and 40.8%, respectively) with a further 23-28% at 0 hpi. Cluster 2 is the only cluster enriched for late infection with 36.7% of its cells from 48 hpi. **Figure 1F** shows the same cluster time point matrix expressed as the percentage of all cells, such that the entire heatmap sums to 100%. This representation highlights which cluster-time point combinations contribute most to the overall population. The cluster 2 population at 48 hpi is the single largest population across the heatmap (8.1% of all cells), whereas the early-enriched clusters (7, 10, and 12) are sparsely populated overall, with no cluster-time point population exceeding 0.6% of total cells.

The representative images in **Figure 1B** show that at the earlier time points of 12 and 20 h the intensity of the E protein signal is above background while the NS4B signal is not. Both channels are clearly detectable from 28 hpi onward. Because DENV is translated as a single polyprotein, E protein and NS4B are produced concurrently, and this heterogeneity therefore reflects the point at which each antigen becomes detectable in our assay rather than the timing of viral protein synthesis.

To further investigate the expression of viral proteins, we trained three XGBoost regression models on the viral channels of the images where 0 and 48 hpi were labeled 0 and 1, respectively: one on both viral channels (NS4B & Epro), one on NS4B only, and one on E protein only. We then scored the probability of infection for cellular objects using each of the three models and averaged scores to the well level. All three models rise from below 0.2 at 0 hpi to approximately 0.78-0.88 at 48 hpi, with strong overlap around 0.5 at 28 hpi (**Figure 2**). The models differ at the earliest time points. When testing the statistical consistency of staining for NS4B and E protein, the E protein model has a statistically significant increase in average well-level score from hour 0 to 12 (p=8.08e-05) and 0 to 20 (p=0.042), whereas the NS4B model does not (p=1 and p=0.824, respectively). Taken together with the representative images in **Figure 1B**, these results are consistent with E protein becoming detectable in this assay before NS4B at early time points of the DENV infection cycle.

**Figure 2:**
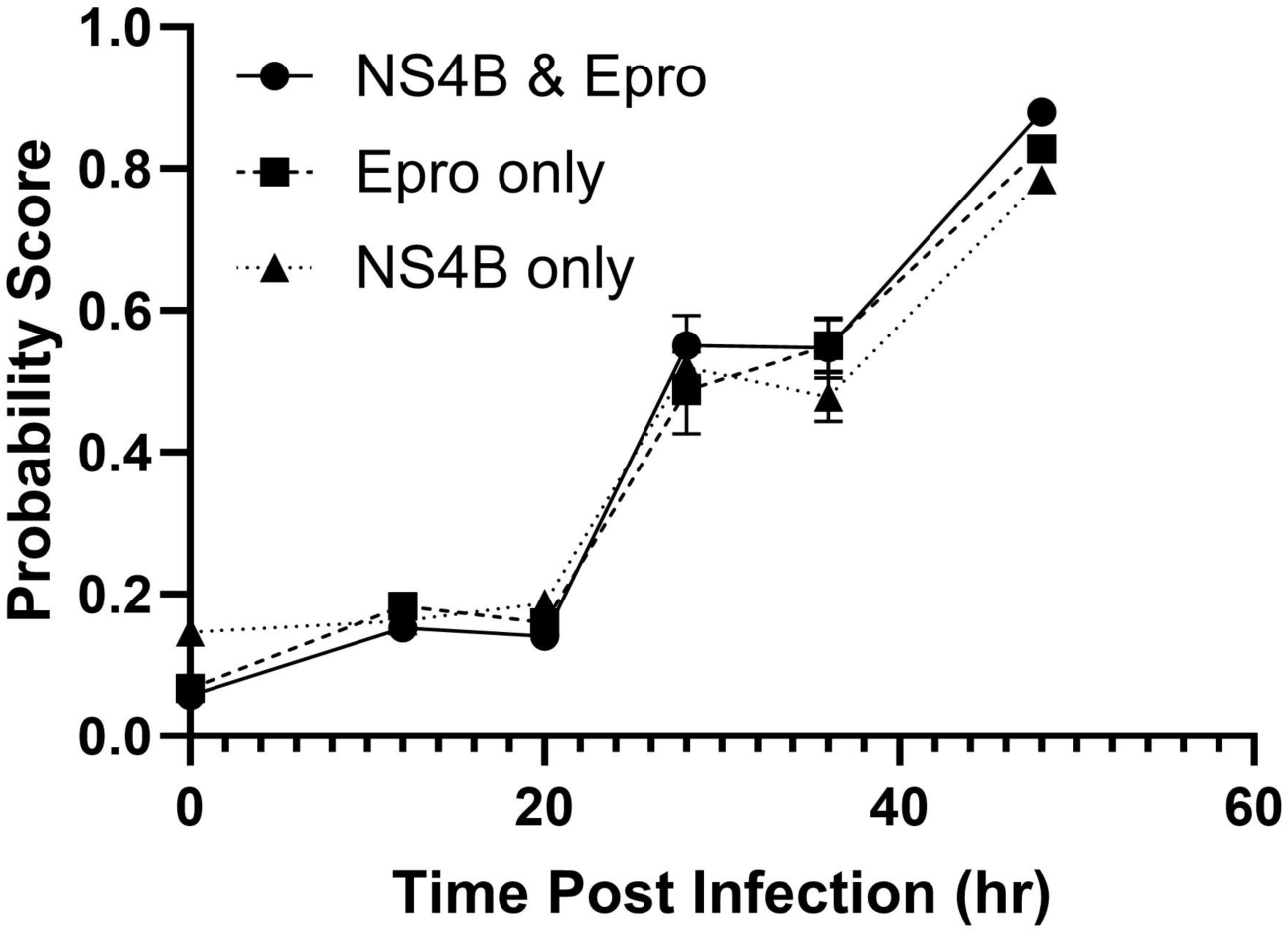
Significant increase in Epro only model probability score at hour 12 shows E protein is detected first at early time points. The prediction of infection probability based on select features increases over time as the infection progresses.

### High-Content Screening to Identify DENV Inhibitors

To identify DENV inhibitors, we screened 960 compounds from the University of Michigan Drug Repurposing Library (FDA-approved drugs and clinical candidates) in a high-content fluorescence imaging assay (384-well format, Huh7.5.1 cells; **Figure 3A**; Methods), using viral protein detection as an infection marker and cell count as a viability readout. Control wells averaged 48.9% infection in 48 hours. The screen identified eight primary hits significantly reducing infection (3-sigma below DMSO controls; hit rate 0.8%, average Z’ 0.7). Seven of these hits, plus two investigational compounds, advanced to concentration-response testing (10-point, 3-fold dilution from 8 μM, 24 h pre-incubation, then DENV-2 infection and 48 h incubation; Methods). Percent infection was quantified via a random forest classifier trained on manually annotated cells (CellProfiler Analyst), and viability from cell counts normalized to DMSO controls. From the concentration-response study, four drugs produced concentration-dependent inhibition of DENV infection relative to infected and mock controls (**Figure 3B**). Esomeprazole had an IC50 of 877 nM (95% CI: 0.453–2.785 µM); pralatrexate 20 nM; and GW4064 290 nM (neither with a reliable confidence interval). LY411575 had an IC50 of 72 nM (95% CI: 0.059–0.16 µM), reducing percent infected per well to almost zero at the highest concentration (8 μM). Concentration-response curves are shown in **Figure 3C** alongside each drug’s structure and a representative image of the 8 μM condition.

**Figure 3.**
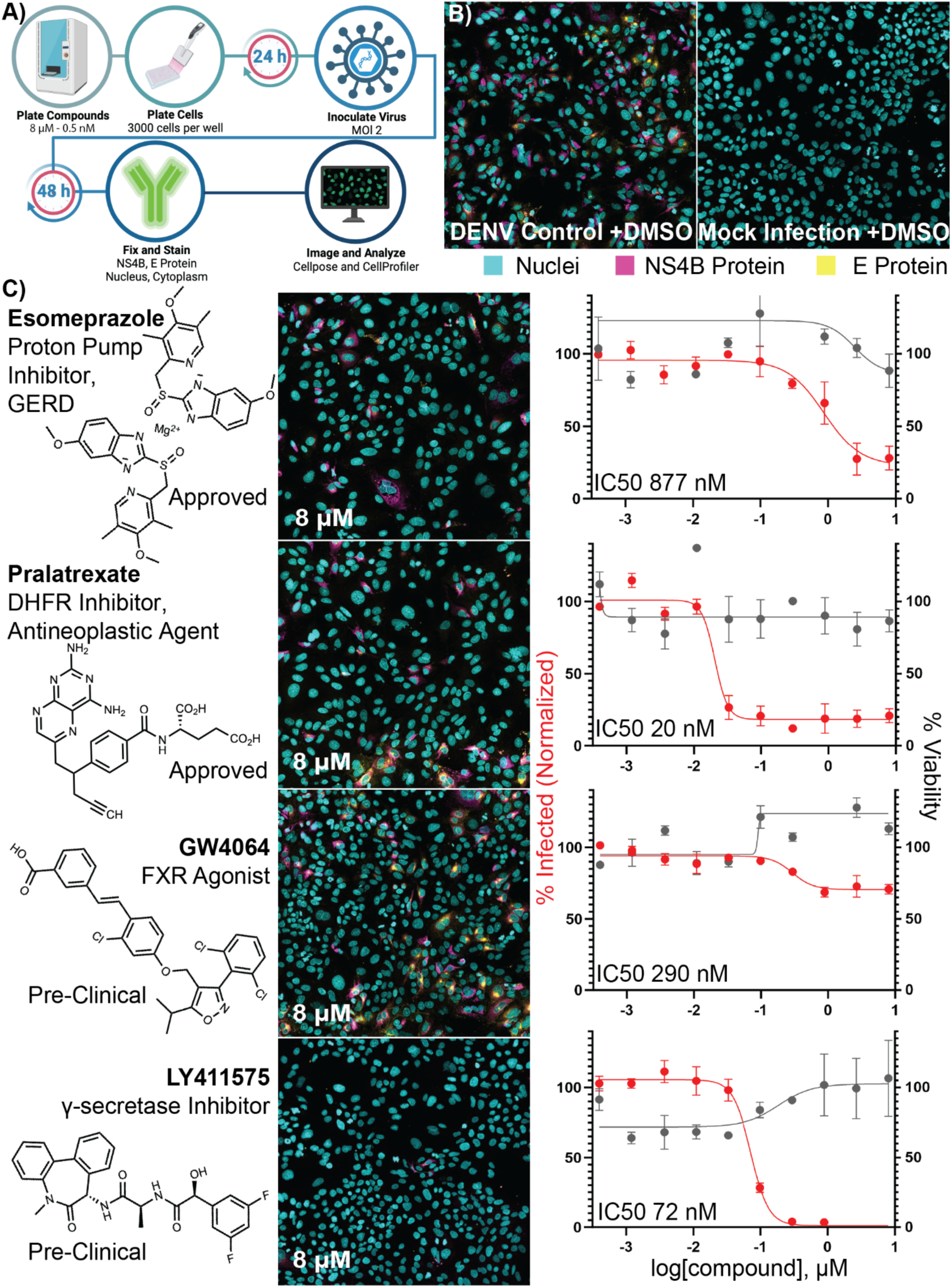
Concentration-Response Data from Drug Repurposing Screen for DENV Antivirals. A.) High-throughput screening workflow. B.) Representative images of the screen controls. C) Drug name, target, and FDA status (left) for each compound in the same row as an image from the highest concentration (center) and the concentration-response curve (right).

### Machine Learning Insights on the Mode of Action for Concentration-Responsive Compounds

With this morphological profiling of the 48-hour DENV life cycle, we used machine learning to predict when a drug arrests infection, training 21 XGBoost regression models on the pseudotime series data and using them to score concentration-response plate images by phenotype. Regressors were trained on all pairwise time point comparisons (e.g., 0 vs. 12) and “one-vs-all” comparisons (e.g., 0 vs. all), so each image carries a 21-dimensional profile of scores describing which infection time points its morphology most resembles.

We embedded these image-level score profiles with UMAP and colored by the predicted time point from a representative regressor, which shows that most conditions across the plate are predicted to resemble late infection, clustering around 48 hpi (**Figure 4A**). A single dense region at the top of the embedding is the only deviation from this, with predicted time points clustering around 12 hpi. Overlaying the LY411575 images onto the same embedding (**Figure 4B**) singled out the same top region: the two highest concentrations (2.7 and 8 μM) of LY411575 occupy it almost exclusively, while lower concentrations distribute throughout the embedding. This concentration-dependent localization indicates that increasing LY411575 exposure shifts morphological phenotypes toward the earliest predicted infection state. Some pralatrexate images also localize to this early-predicted region (**Supplementary Figure S3**), but its distribution remains largely diffuse and is notably adjacent to and distinct from the LY411575 region which extends furthest into the earliest predicted phenotypic space.

**Figure 4.**
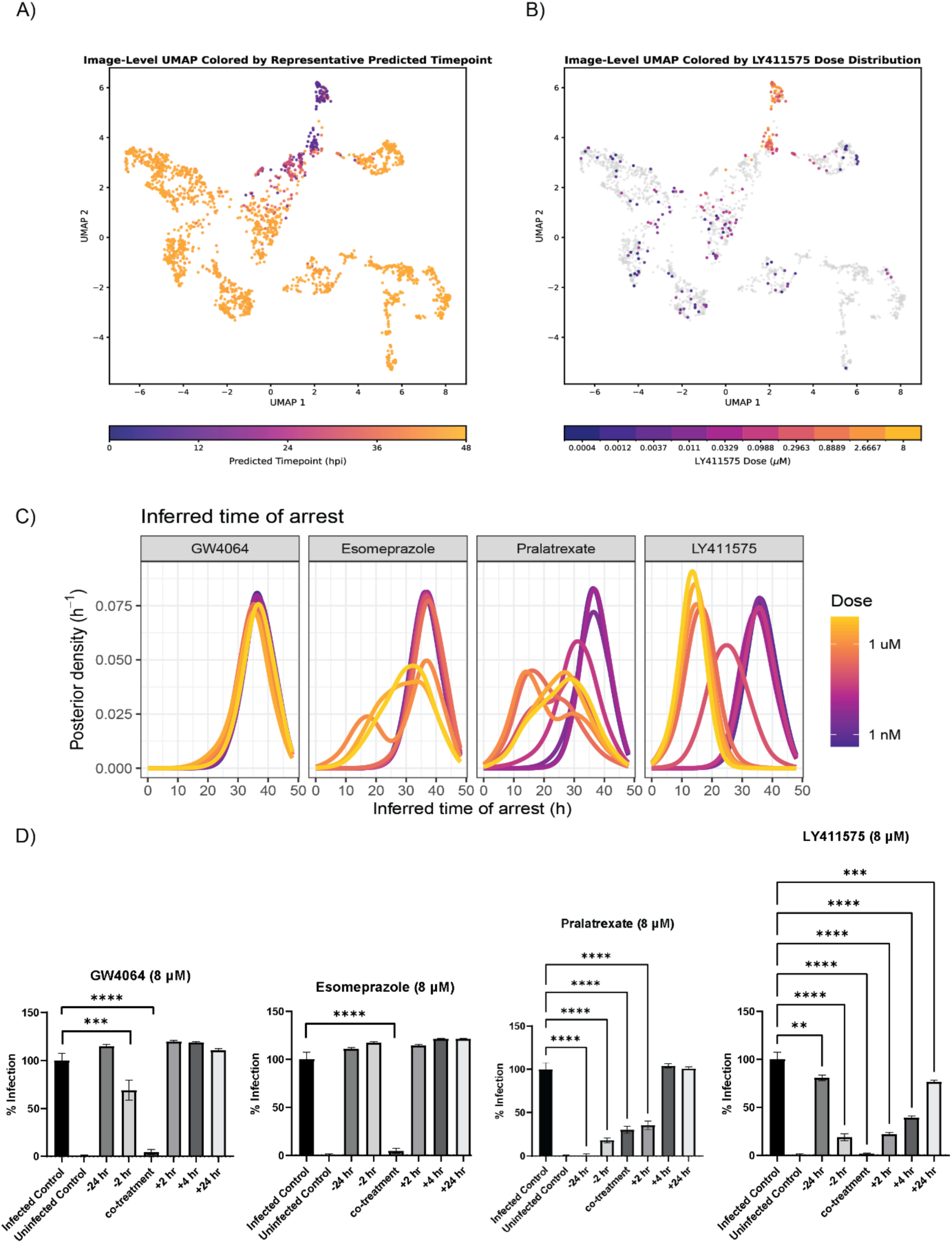
Mapping antiviral phenotypes to infection-time space using machine-learning regression derived predictions. **A.)** Image-level UMAP of all concentration-response conditions colored by predicted time point using the all vs 48 hpi XGBoost regressor model as a representative model. B.) Image-level UMAP from A.) highlighting the dose distribution of hit compound LY411575. C.) The inferred time of arrest across all drugs and doses, estimated by the posterior of a time point under a mixture of Student’s *t* likelihood distributions weighted by the XGBoost model scores and a uniform prior. D.) Time of addition experiment results with repurchased compounds. Statistical comparisons are to the infected control within each compound (Dunnett’s test). *P < 0.05, **P < 0.01, ***P < 0.001, ****P < 0.0001. n = 24 (infected/uninfected control), n = 14 (all time-of-addition conditions).

Using these models, we estimated the likelihood of a drug arresting infection at an earlier time point, reasoning that a high score under a discriminative model indicates arrest near that time point. We defined Student’s *t*-distributions around each of the 21 XGBoost models’ conditions and, given the observed scores for the four drug-treated cell morphologies (**Supplementary Figure S4**), estimated the posterior across 0-48 h (**Figure 4C**). For all compounds, the mean and standard error over three well-replicates of the posterior mean at zero concentration ranged between 35.7-36.2 [± 0.03-0.28]. For GW4064 at the max concentration of 8 μM, the estimated time of arrest was 35.9 [± 0.1] h, indicating no significant change. Esomeprazole’s estimated time of arrest at 8 μM was 29.9 [± 3.4] h, showing a modest and more variable effect, while LY411575 and pralatrexate showed distinct, concentration-dependent predictions: Pralatrexate’s estimated time of arrest had a minima at 0.28 μM of 21.4 [± 3.2] h and then rose to 26.1 [± 4.0] h at 8 μM. LY411575 monotonically shifted cellular phenotypes to early infection with a dose of 0.1 μM giving 26.2 [± 1.3] h doses of 8 μM giving 13.8 [± 0.2] h, reflecting a concentration-dependent effect to an early time of infection. Posterior mean time of arrest estimates are shown for all nine compounds tested in concentration-response in **Supplementary Figure S5.**

To test the time of arrest estimates, we performed a time-of-addition experiment with the four repurchased compounds, adding each at 8 μM at time points from 24 h before infection to 24 h after and measuring infection (**Figure 4D**). Esomeprazole and GW4064 inhibited infection only under co-treatment, with no significant reduction at any post-infection addition time, illustrating both compounds are active only at or immediately around viral entry. This agrees with the likelihood analysis in which neither compound produced strong evidence of arresting infection before 36 hpi.

LY411575 and pralatrexate showed the opposite profile: LY411575 significantly reduced infection at every time tested from −24 h to +24 h, with complete inhibition under co-treatment, and pralatrexate inhibited infection from −24 h to +2 h, with activity lost by +4 h. Unlike the entry-restricted profiles of esomeprazole and GW4064, both inhibit DENV when added after inoculation, indicating a post-entry step. Because retained post-inoculation activity argues against a pure entry block, we interpret the model’s output as the point at which the infected population is morphologically arrested rather than the specific viral process inhibited, and the time-of-addition results broadly support the pseudotime model’s mode of action assignments.

## Discussion

There remains an urgent need for DENV therapeutics, which can be used effectively to reduce the burden of severe dengue on developing countries’ health infrastructure, especially due to complications with the vaccine. Herein we report a streamlined computational workflow that provides insight into high-throughput drug screen results to expedite drug discovery.

A notable feature of our DENV pseudotime infection model is that E protein becomes detectable before NS4B (**Figure 1B, 2**). We do not interpret this as a difference in viral protein synthesis, since all DENV proteins are translated from a single polyprotein precursor and E/NS4B expression is intrinsically coupled prior to proteolytic processing.^31^ We instead attribute the temporal asymmetry to assay detection, possibly reflecting antibody affinity/epitope accessibility or subcellular distribution differences: E protein accumulates in dense deposits at the ER and plasma membrane,^2^ whereas NS4B is distributed across replication-complex membranes, so the two stains cross the assay’s detection threshold at different points in infection. We use this asymmetry to discriminate early time points that would otherwise be indistinguishable in a single-channel assay, positioning compound-treated wells along the infection course for mode of action inference.

From the high-content screen, the FDA-approved drugs esomeprazole and pralatrexate and the commercially available GW4064 and LY411575 showed potent antiviral activity against DENV (**Figure 3**). Esomeprazole, a proton-pump inhibitor used for GERD and gastric acid hypersecretion, has shown antiviral activity beyond its approved indication: it was effective against SARS-CoV-2 delta.^32^ Additionally, both esomeprazole and the related PPI tenatoprazole have been shown to inhibit HIV-1 replication by covalently modifying the ESCRT-I protein Tsg101, disrupting its ubiquitin-binding activity, and preventing recruitment of Tsg101 to the plasma membrane viral budding site, thereby impairing viral egress.^33^ In this same study, tenatoprazole showed no activity against DENV-2 produced in 293T cells at up to 100 μM when scored by focus-forming assay on Vero monolayers. However, esomeprazole itself was not tested, so our result is not in direct conflict with that report.^33^ Given other viruses analogously exploit Tsg101, including alphacoronaviruses^34^ and flaviviruses, this pathway represents a plausible mechanism underlying the antiviral activity observed here.

Pralatrexate is a DHFR inhibitor with satisfactory bioavailability used to treat relapsed or refractory peripheral T-cell lymphoma. In a drug-repurposing screen, pralatrexate was found to inhibit SARS-CoV-2, but the authors did not describe a mechanism of action.^35^ DHFR inhibitors are not specific to viral targets,^36^ but there was no significant reduction in cell viability at the concentrations we tested indicating a significant cell-based therapeutic index.

GW4064 was first described in 2000 as a farnesoid X receptor (FXR) agonist later discovered to have poor bioavailability (10%).^37,38^ GW4064 was found to stimulate FXR, a bile acid receptor, and disrupt lipid homeostasis in rotaviruses, which the authors propose could also be a valid mechanism to inhibit DENV replication, as cellular lipids play multiple roles in the orthoflavivirus infection cycle.^39^ Due to GW4064’s poor bioavailability, other FXR agonists could be further explored for broad antiviral activity.

LY411575, an antagonist of the catalytic domain of gamma-secretase (presenilin), has been shown to inhibit HCV, also in the Flaviviridae family,^40^ by inhibiting the presenilin-like aspartic protease signal peptide peptidase (SPP), preventing cleavage of the core protein’s C-terminal transmembrane region during maturation.^40^ A recently reported druggable genome CRISPR screen identified SPP as a host factor for DENV infection and also found that LY411575 effectively arrests DENV replication.^41,42^ No experimental data are available, to our knowledge, on LY411575’s oral bioavailability.

Given that raw regression scores do not directly correspond to a time point, scores were transformed into weighted probabilities via a Student’s *t*-distribution, enabling prediction of each compound’s ability to arrest infection along a continuous DENV temporal trajectory. The resulting posterior estimates revealed distinct compound-specific patterns that provided insight into their potential antiviral modes of action (**Figure 4C**). Lower concentrations of all four hits predicted arrest near one of the latest time points (36 hpi), consistent with inadequate suppression, while LY411575 exhibited the clearest concentration-dependent shift toward earlier stages. These predictions suggest that LY411575 induces a phenotype resembling early infection at high concentrations, effectively arresting the virus and preventing progression to a later, more mature phenotype. Consistent with the concentration-dependent shift in inferred time of arrest, UMAP analysis showed that samples treated with the highest concentrations of LY411575 clustered within a distinct region of phenotypic space (**Figure 4B**).

Importantly, time-of-addition experiments were broadly consistent with these computational predictions. LY411575 significantly reduced infection when administered before, during, and notably after inoculation, with the strongest effects at −2 h, co-treatment, and +2 h (**Figure 4D**). This would be consistent with previous reports identifying SPP as the molecular target of LY411575 and implicating SPP in flaviviral replication.^42^ Qiao and colleagues recently reported LY411575 and other SPP-targeting compounds as broad-spectrum flavivirus inhibitors via this mechanism, with substantially higher potency in the same cell line (EC50 3-5 nM) than we observe here (IC50 72 nM), a gap likely reflecting assay or readout differences between studies.^42^ Although our data do not conclusively establish a viral mechanism, the convergence of computational and experimental findings positions LY411575 as the most compelling candidate for mechanistic follow-up studies.

Pralatrexate produced a broader and more heterogenous posterior at higher concentrations, with no single dominant inferred time of arrest (**Figure 4C**), which may reflect a phenotype outside the training data’s range. Notably, time t=0 h in our pseudotime model corresponds to mock infected cells rather than drug-treated cells. Therefore, compounds that strongly perturb host-cell morphology while preventing viral infection would not necessarily be expected to resemble the t=0 h reference state. Instead, such perturbations may generate phenotypes that fall outside the pseudotime infection continuum used for model training, resulting in a diffuse posterior across inferred times of arrest. Independently, unsupervised clustering showed a more dispersed phenotypic distribution for pralatrexate across all concentrations (**Supplementary Figure S3**), consistent with a phenotypic signature not well described by any single trained time point. Time-of-addition experiments showed maximal activity when pralatrexate was added prior to infection (−24 h, −2 h), with activity progressively lost at later times. Together, these observations suggest an early-stage antiviral effect and illustrate how uncertainty within the pseudotime framework can itself be mechanistically informative.

GW4064 and esomeprazole, by contrast, showed limited to no concentration-dependent shifts in inferred arrest time (**Figure 4C**), consistent with their co-treatment-restricted activity window (**Figure 4D**). For esomeprazole, the near-complete suppression observed during co-treatment compared to other conditions suggests the antiviral activity is confined to a narrow temporal window, which may indicate an entry or immediate post-entry mode of action. Wu and co-workers suggested GW4064 indirectly downregulates a receptor required for HCV entry,^43^ raising the possibility that GW4064 similarly acts as a flavivirus entry-inhibitor, though the role of FXR in flavivirus replication needs further study.

Together, these results show that combining multiple regression predictions can resolve compound-specific phenotypic shifts that map onto DENV’s temporal dynamics, distinguishing potential entry and later-stage infection inhibitors and generating testable mode of action hypotheses from imaging data alone. Further, the uncertainty of the likelihood analysis may be biologically informative itself, suggesting that these cells occupy a phenotypic state that is not captured during model training. Collectively, this study demonstrates that image-derived phenotypes can be leveraged not only to identify antiviral compounds, but to also infer their position on the DENV infection trajectory.

This study has several limitations. Pseudotime and concentration-response experiments were performed on a single plate with time points arranged in blocks rather than being randomized, therefore time-associated effects cannot be fully separated from potential plate-position effects. Target elucidation of our hits is necessary to confirm mechanistic predictions, which are derived from two viral protein markers. Time-of-addition profiling used the same image-based readout as the other DENV bioassays and therefore does not constitute orthogonal confirmation of antiviral activity. All experiments were conducted in a single cell line (Huh7.5.1) with a single DENV serotype; generalization to other cell types and serotypes is necessary to translate the findings presented here.

We identified four potential antivirals for DENV and developed a likelihood-based framework that predicts the time of arrest of infected cells using image-derived morphological features. The identification of LY411575, previously characterized as a validated DENV antiviral, supports our framework’s ability to recover biologically relevant signatures. Similar image-based approaches have mapped infection stages and dynamics for other viruses^44–48^, but our probabilistic scoring framework generating a continuous, concentration-related estimate of infection timing is, to our knowledge, novel, offering a scalable means of generating temporal hypotheses from high-content screens for antiviral discovery and mechanistic prioritization. In future work, we plan to extend this approach to other viruses and pursue target validation of the hypotheses generated here.

## Author Contribution

Conceptualization: J.G.H, J.Z.S., M.J.O.

Data Curation: J.G.H., S.F.B, B.H., J.Z.S, M.J.O.

Formal Analysis: J.G.H., S.F.B., J.Z.S., M.J.O.

Funding acquisition: J.Z.S., M.J.O.

Investigation: J.G.H., S.F.B., J.W.W, S.P., D.M.M.J., A.W.T., J.Z.S, M.J.O.

Methodology: J.G.H., S.F.B., J.W.W, S.P., D.M.M.J., A.W.T., J.Z.S, M.J.O.

Project administration: J.Z.S, M.J.O.

Resources: A.W.T., J.Z.S.

Software: J.G.H., S.F.B, B.H., J.Z.S, M.J.O.

Supervision: A.W.T., J.Z.S, M.J.O.

Validation: J.G.H., S.F.B., J.W.W, S.P., D.M.M.J., A.W.T., J.Z.S, M.J.O.

Visualization: J.G.H., S.F.B., J.W.W, S.P., D.M.M.J., A.W.T., J.Z.S, M.J.O.

Writing - original draft: J.G.H, S.F.B.,

Writing - review & editing: J.G.H., S.F.B., J.W.W, S.P., B.H., D.M.M.J., A.W.T., J.Z.S, M.J.O.

## Acknowledgements

The authors acknowledge the use of the Yokogawa Cell Voyager 8000 in the University of Michigan BRCF Microscopy Core, purchased with support from NIH S10 grant 1S10OD034245.

## Data, Metadata, and Code Availability

Raw high-content image data and associated image analysis pipelines are available through Zenodo at https://doi.org/10.5281/zenodo.22877652. All code used for data processing, analysis, and visualization, along with associated analysis files, is available in the GitHub repository at https://github.com/maomlab/DenguePseudoTime.

---

**Supplementary Figure S1.**
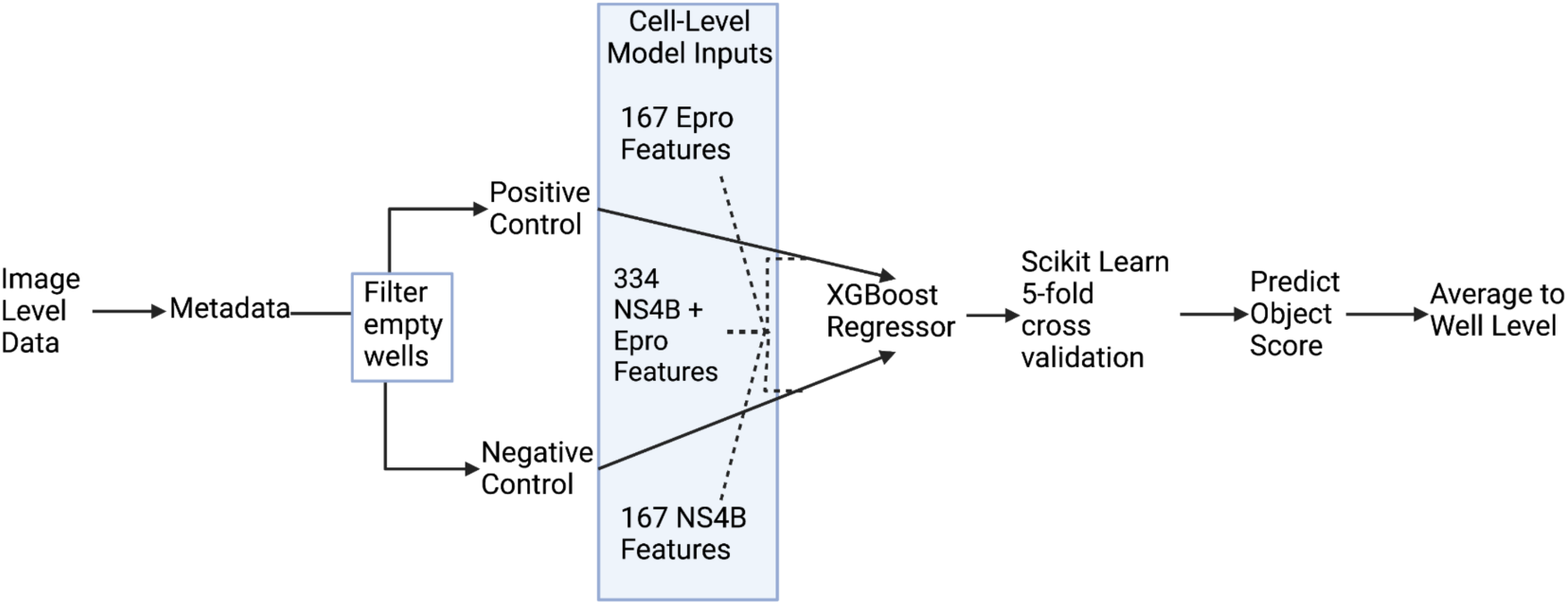
Machine Learning Model Workflow.

**Supplementary Table S1.** R^2^ Values for Drug Treatment vs. Time Point Models.

|  | 0 h | 12 h | 20 h | 28 h | 36 h | 48 h |
| --- | --- | --- | --- | --- | --- | --- |
| 0 h |  | 0.741 | 0.748 | 0.936 | 0.928 | 0.891 |
| 12 h |  |  | 0.654 | 0.941 | 0.941 | 0.950 |
| 20 h |  |  |  | 0.735 | 0.820 | 0.886 |
| 28 h |  |  |  |  | 0.516 | 0.847 |
| 36 h |  |  |  |  |  | 0.677 |
| all | 0.758 | 0.626 | 0.375 | 0.623 | 0.496 | 0.831 |

**Supplementary Figure S2:**
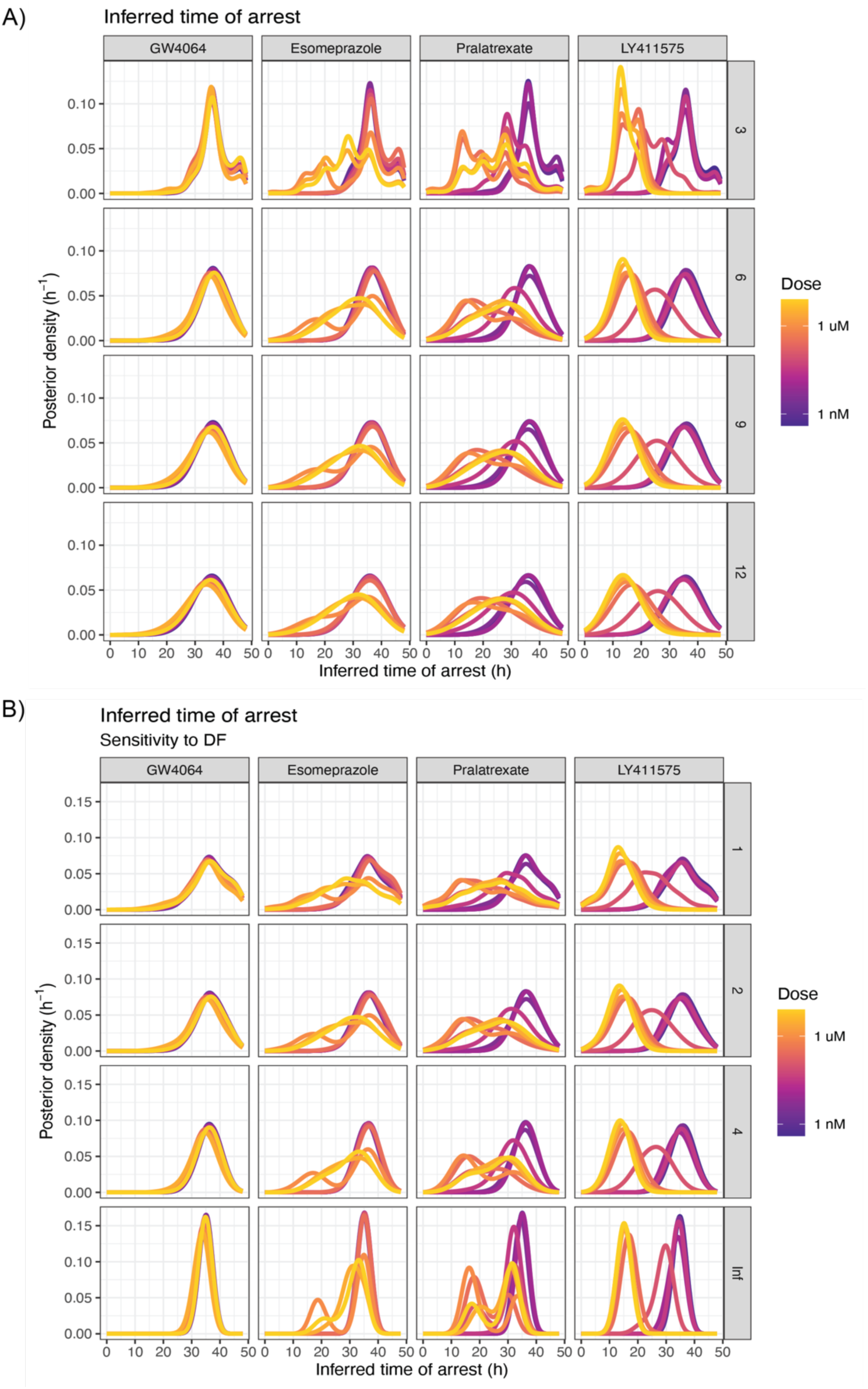
Sensitivity of inferred time of arrest posterior densities to Student’s *t*-distribution parameters. Curves are colored by dose. A.) Sensitivity to the scale, *σ*, of the Student’s *t*-distribution across scale values 3, 6, 9, and 12. B.) Sensitivity to the degrees of freedom (df), *ν*, of the Student’s *t*-distribution across df = 1, 2, 4 and ∞.

**Supplementary Figure S3:**
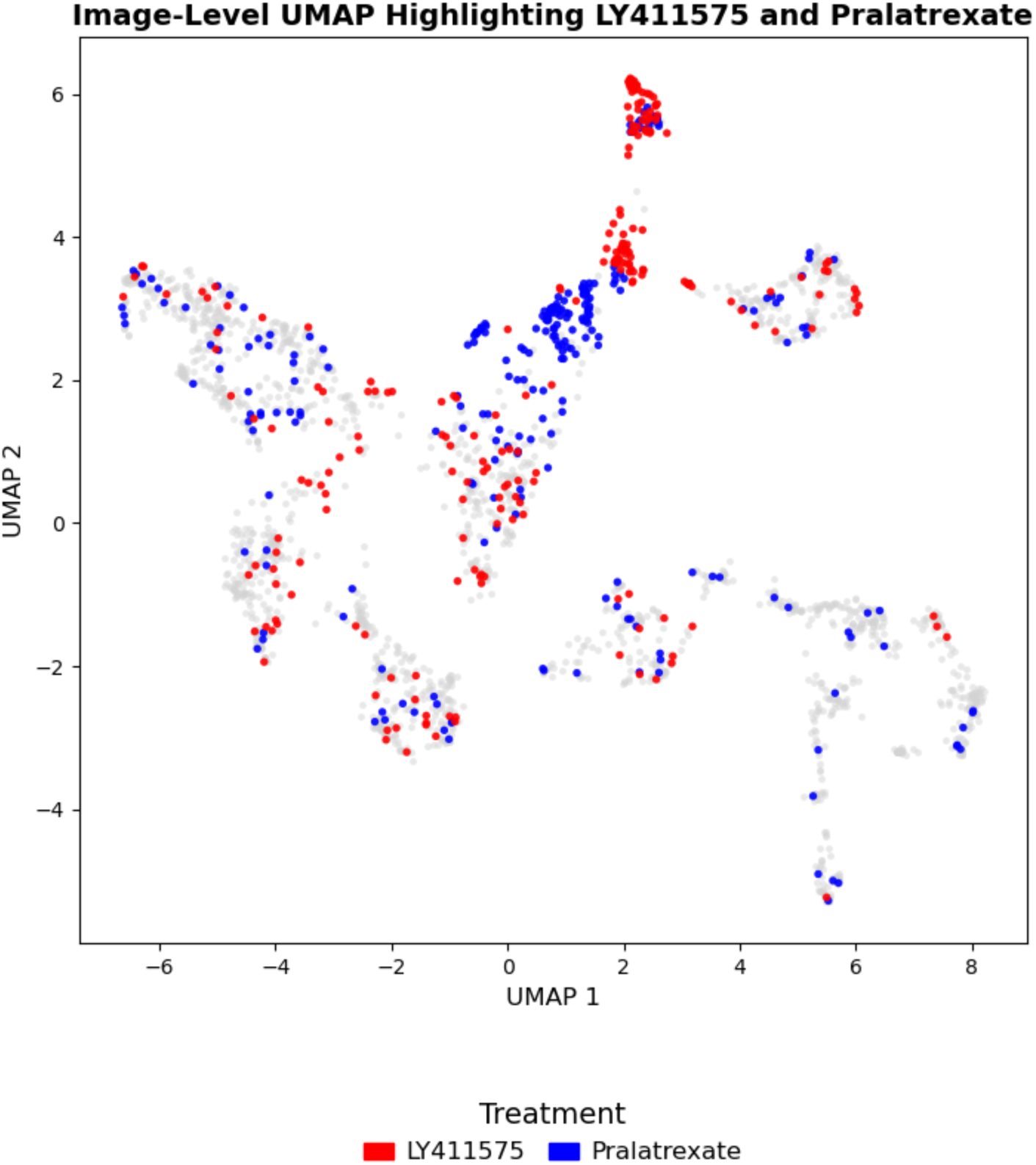
Image-level UMAP highlighting LY411575 (red) and pralatrexate (blue) across all dose-response experiment wells. Grey points represent all other wells. Both compounds preferentially localize to the region of the embedding predicted to closely resemble early infection, with high-concentration LY411575 wells extending to the uppermost region.

**Supplementary Figure S4:**
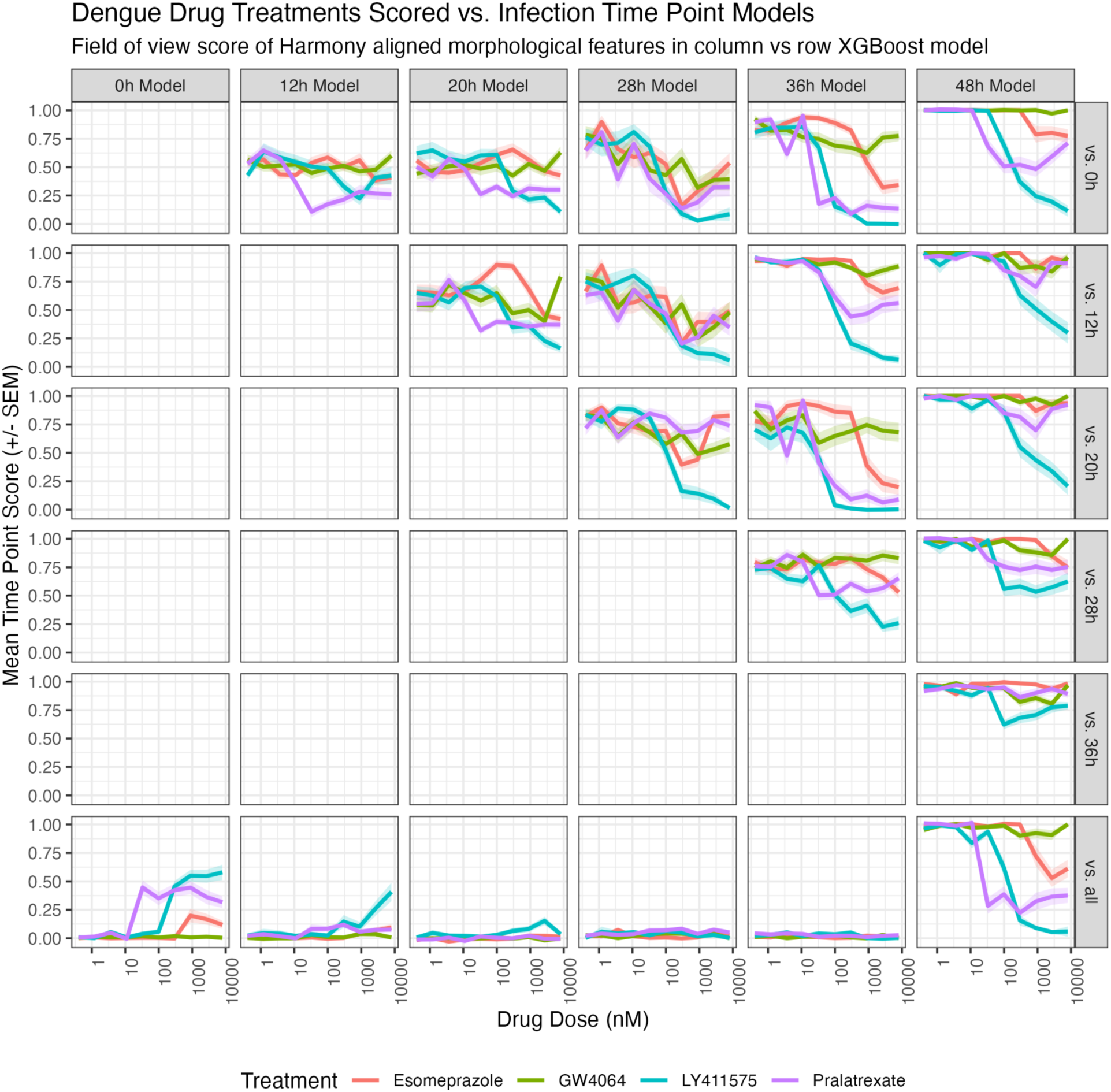
Scores of drug-treatment concentration-response morphologies under 21 XGBoost models trained to discriminate the column vs row time points (above the diagonal) or trained to discriminate the column time point vs all other time points (bottom row). Scores are represented as the mean score ± standard error in the mean across fields of view.

**Supplementary Figure S5:**
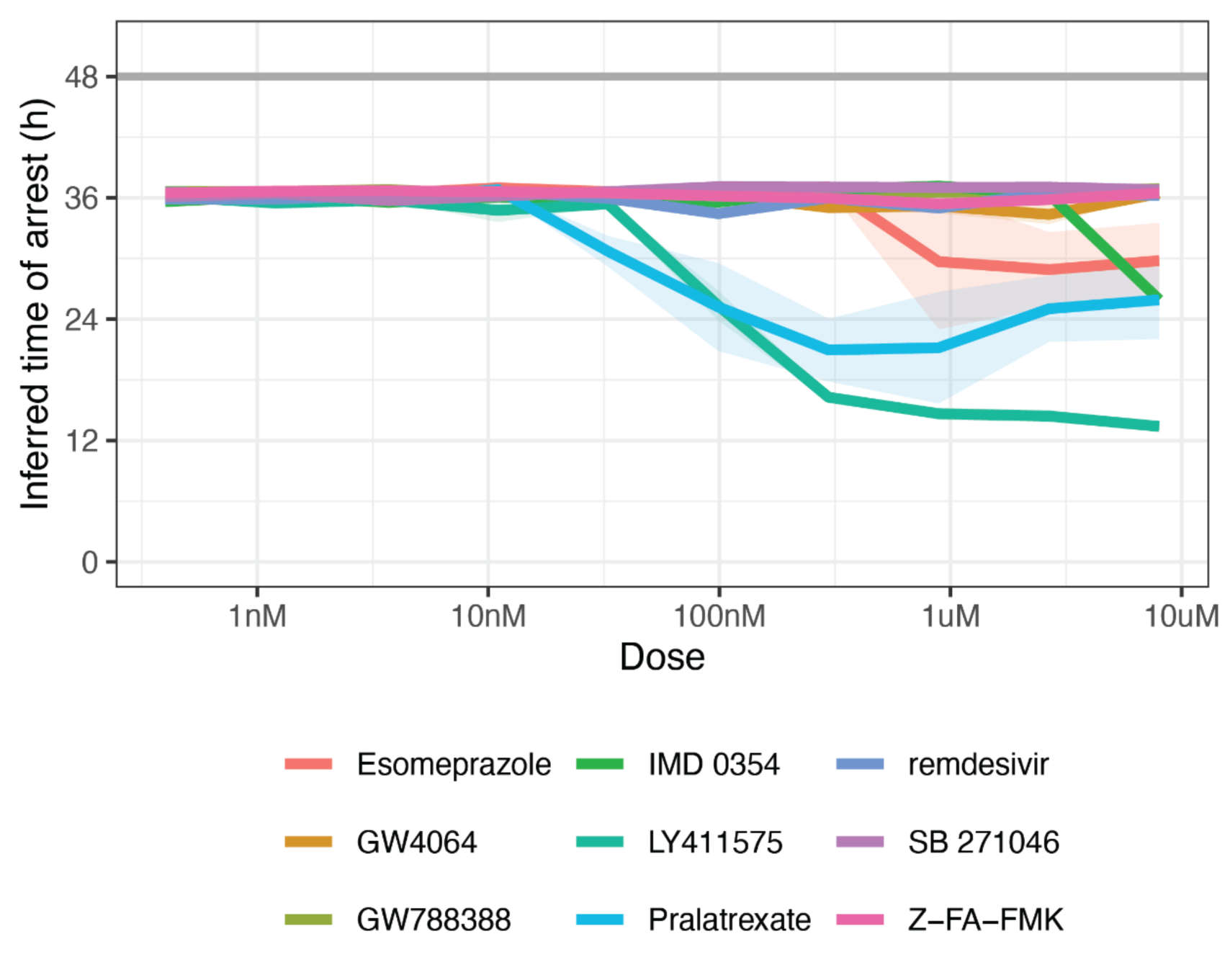
Concentration-dependent changes in inferred time of arrest across all nine compounds tested in concentration-response experiments. The posterior mean inferred time of arrest across three well-replicates is plotted as a function of compound concentration ± standard error in the mean across well-replicates.

